## Supplemental Table S5 for "Yeast NDI1 reconfigures neuronal metabolism and prevents the unfolded protein response in mitochondrial complex I deficiency"

| **Stock** | **Source & stock number** | **Construct ID** |
| --- | --- | --- |
| *w^1118^* | BDSC 6326 |  |
| *nSyb-Gal4* | From Rita Sousa-Nunes, King’s College London |  |
| *UAS-Dcr2;OK371-Gal4,UAS-CD8-GFP* | From Darren Williams, King’s College London |  |
| *y^1^,v^1^;UAS-PERK dsRNA* | BDSC 42499 | HMJ02063 |
| *Daughterless GeneSwitch-GAL4* | Nazif Alic, University College London |  |
| *Tubulin-GAL80^ts^* | BDSC 7108 |  |
| *UAS-ND-75 RNAi* | BDSC 33910 | HMS00853 |
| *UAS-ND-75 RNAi* | BDSC 33911 | HMS00854 |
| *UAS-mitoGFP* | BDSC 8442 |  |
| *UAS-SPLICS_L_* | This study |  |
| *UAS-IP_3_R RNAi* | BDSC 51686 | GLC01786 |
| *UAS-Hsc70-3^K97S^* | BDSC 5842 |  |
| *UAS-Hsc70-3^D231S^* | BDSC 5841 |  |
| *UAS-NDII* | Alex Whitworth, MRC Mitochondrial Biology Unit, Cambridge (Sanz et al., 2010) |  |

10.1073/pnas.0911539107.
